# *KLF17* acts as a dual reporter for tracking naïve pluripotency and 8-cell-like states

**DOI:** 10.64898/2026.10.23.761375

**Authors:** Sıla Unlu, Arda Odabas, Nilya Karasurmeli, Tamer T. Önder

## Abstract

Human pluripotent stem cells (hPSCs) can be stabilized in states corresponding to successive stages of preimplantation development. Conventional primed hPSCs resemble the post-implantation epiblast, naïve hPSCs the pre-implantation epiblast, and rare 8-cell-like cells (8CLCs) arising within naïve cultures recapitulate features of zygotic genome activation (ZGA) at the 8-cell stage of early human embryonic development. To enable monitoring of these successive states, we exploited the expression of *KLF17*, which is absent from primed hPSCs and present in the human 8-cell embryo at levels exceeding those of the epiblast. We hypothesized that its expression level, rather than its presence alone, would report position along the continuum of early human development. We therefore generated an EGFP knock-in at the endogenous *KLF17* locus and asked whether expression level could resolve two consecutive transitions in the same line. During conversion of primed cells to the naïve state, KLF17-positive cells showed elevated expression of naïve-associated genes including *SUSD2, DNMT3L, DPPA3, DPPA5, TFCP2L1* and *CD75* relative to the negative fraction, and formed blastoids at higher efficiency. Higher *KLF17* levels in turn marked the naïve-to-8C-like transition: following TSA and DZNep treatment, cells sorted into negative, low, mid and high fractions showed a progressive increase in ZGA-associated transcripts such as *ZSCAN4*, *LEUTX, TPRX1, MBD3L2, DUXA* and *RFPL2. KLF17* expression level thus provides a single continuous readout of position along early human development, from the primed state through naïve identity to the 8C-like state.

## Introduction

Pluripotency is a transient property of the human embryo, present for roughly ten days between the emergence of the epiblast in the blastocyst and lineage commitment at gastrulation^1^. *In vitro*, this window has been captured in more than one form. Conventional human pluripotent stem cells (hPSCs), whether derived from blastocysts^2,3^ or generated by somatic cell reprogramming^4^, correspond transcriptionally to the post-implantation, gastrulating epiblast^5^ and are referred to as primed. Cells resembling the pre-implantation epiblast, naïve hPSCs were described more recently and can be obtained by resetting of conventional hPSCs^6–8^, by direct derivation from dissociated day-6 blastocysts^9^, or during somatic cell reprogramming^10,11^. The two states differ in transcription factor networks, global DNA methylation, X chromosome status, metabolic profile and colony morphology and signaling requirements^6,7,8,12^. Conversion of hPSCs between these states in vitro offers an attractive model to study early cell fate decisions.

The developmental range accessible in culture has since been extended to earlier stages. Cells transcriptionally resembling the 8-cell embryo were first observed as a transient population in naïve cultures^13–15^ and were termed 8CLCs. Taubenschmid-Stowers et al. defined this program and found that ∼1% of naïve cells were marked by high *TPRX1* expression, and 8-cell-specific genes^13^. Mazid et al. increased the 8CLC fraction to ∼13% through epigenetic modulation via TSA and DZNep treatment and used a *TPRX1* reporter to isolate cells with transcriptional and epigenetic features of the human 8-cell embryo^14^. Yu et al. also independently confirmed 8CLCs by using a *LEUTX*-based reporter system^15^. More recently we have described a chemical induction cocktail that can induce up to 40% 8CLCs within 48h from naïve cultures^16^.

These states were discovered by transcriptomic profiling but could only be studied extensively once a reporter enabled their isolation from heterogeneous cultures. Naïve identity is typically assessed with cell-surface markers such as SUSD2, CD75, CD7 and CD130^17,18^ or with genetically encoded reporters such as ΔPE-OCT4-EGFP, which exploits the fact that *POU5F1* is transcribed from different enhancers in the two states^7^. On the other hand, 8CLCs are assessed with TPRX1, LEUTX-based reporters or by the expression of cell surface receptor SLC34A2^19^. Because each reporter captures only one transition, following progression from naïve to 8C-like identity requires an additional reporter, and therefore a second engineered allele or fluorescent marker.

KLF17 is a naïve-specific transcription factor, essentially absent from primed hPSCs and used as a defining marker of the pre-implantation epiblast ^18,20^ but it is also expressed in the human 8-cell embryo, where its levels exceed those seen in the epiblast. We therefore asked whether a KLF17-based reporter can track both acquisition of naïve pluripotency and progression toward an 8-cell-like state. We generated a EGFP knock-in at the endogenous KLF17 locus and found that reporter intensity distinguished naïve from primed cells and increased with expression of ZGA-associated genes upon transition to 8CLCs. KLF17 loss impaired naïve reprogramming, while reporter-positive naïve cells formed blastoids more efficiently. Thus, a single KLF17-EGFP reporter provides a graded readout across the primed-to-naïve and naïve-to-8C-like transitions.

## RESULTS

### *KLF17* is induced in two distinct phases during human preimplantation development and marks the naïve state *in vitro*

To determine whether *KLF17* expression marks more than one cellular state, we first examined its dynamics across human preimplantation development. In the single-cell reference map of human embryogenesis^21^, *KLF17* expression was essentially absent at the zygote and 2–4-cell stages, sharply induced at the 8-cell stage, and declined thereafter through the morula and early ICM (Fig. 1A, B). In the late blastocyst, *KLF17* is undetectable in trophectoderm and primitive endoderm but is re-expressed in the epiblast, producing a biphasic pattern with an early peak coinciding with zygotic genome activation and a later, lineage-restricted phase in the pluripotent compartment. Quantification of published scRNA-seq data^22^ supported this observation (Fig. 1C). *KLF17* was highest at the 8-cell stage, fell progressively through morula and early ICM, and was negligible in primitive endoderm, but reached levels in the epiblast comparable to those of the morula. Thus, *KLF17* expression shows an early peak at the 8-cell stage, declines through the morula and early ICM, and rises again in the epiblast.

**Figure 1.**
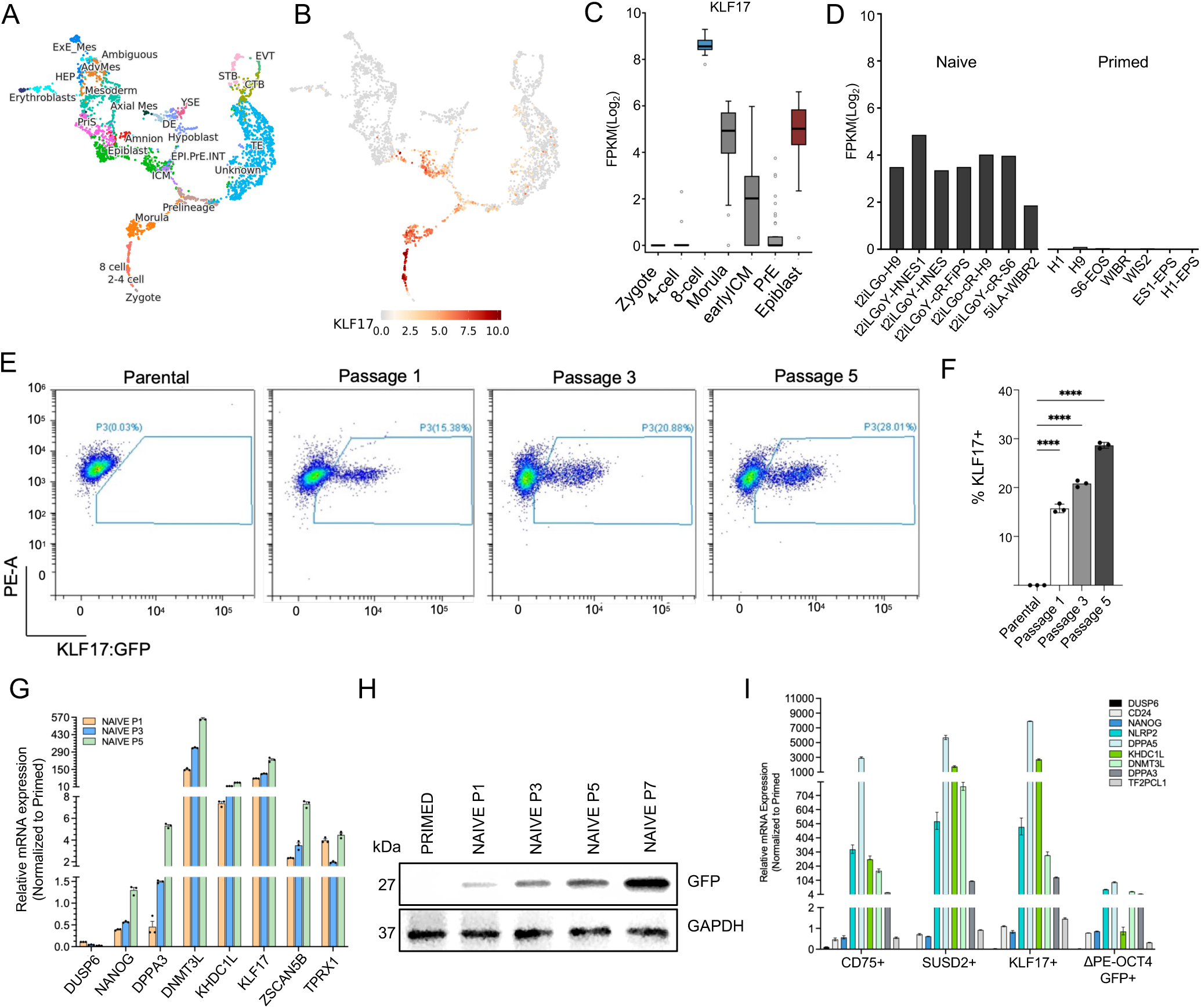
KLF17 is induced in two distinct phases during human preimplantation development and marks the naïve state in vitro. A) UMAP projection of single-cell RNA-seq data from human preimplantation embryos (Petropoulos et al., 2016), annotated by developmental stage and lineage. B) *KLF17* expression projected onto the same UMAP. Color scale indicates normalized expression. C) *KLF17* expression across staged embryonic populations, quantified from published RNA-seq data (Stirparo et al., 2018). EPI, epiblast; PrE, primitive endoderm. D) *KLF17* expression in publicly available RNA-seq datasets across naïve and primed human ES and iPS cell lines. E) Flow cytometry of *KLF17*:EGFP in the untargeted parental line and in the reporter line at passages 1, 3 and 5 of primed-to-naïve conversion. F, Quantification of E. Bars show mean ± SD of 3 independent experiments; individual points are shown. ****p < 0.0001. G) RT-qPCR for primed (*DUSP6*), naïve (*NANOG, DPPA3, DNMT3L, KHDC1L, KLF17*) and 8C-associated (*ZSCAN5B*, *TPRX1*) genes at passages 1, 3 and 5 of naïve conversion. Expression normalized to GAPDH and shown relative to primed. Mean ± SD of 3 biological replicates. H) Immunoblot for EGFP and GAPDH in primed K2G reporter cells and at passages 1, 3, 5 and 7 of naïve conversion. I) RT-qPCR for primed- and naïve -associated genes in CD75-positive, SUSD2-positive, *KLF17*-EGFP-positive and ΔPE-OCT4-EGFP-positive fractions sorted in parallel from the passage-2 conversion culture. Expression normalized to GAPDH and shown relative to primed.

We next examined whether this distinction is preserved in cultured hPSCs. Across a panel of naïve and primed human ES and iPSC lines, *KLF17* was expressed in all naïve lines examined, whereas expression was at or near background in every primed line (Fig. 1D). These observations led us to hypothesize that the level of *KLF17*, rather than its presence alone, could discriminate three configurations within a single line: absent in primed cells, intermediate in naïve cells, and high in 8C-like cells. To test this, we generated a KLF17-EGFP knock-in reporter iPSC line by inserting a T2A-EGFP cassette immediately upstream of the *KLF17* stop codon, together with a LoxP-flanked PGK-Puro selection cassette (Fig. S1A). Correct targeting was confirmed by PCR across both junctions and by amplification of the EGFP and Puro cassettes, while a backbone-specific PCR excluded random integration of the donor plasmid (Fig. S1B). The edited locus was validated by Sanger sequencing (Fig. S1C), and karyotype analysis confirmed that the engineered line had no detectable chromosomal abnormalities following Cas9-mediated targeting (Fig. S1D). This reporter line, which we term *K2G*, retained normal primed and naïve colony morphology (Fig. S1E).

Because primed-to-naïve conversion is asynchronous, we used the reporter to follow the emergence of naïve cells during conversion. Primed K2G cells were transferred to naïve conditions in HENSM^12^ and analyzed by flow cytometry at successive passages, with the parental line included as a gating control (Fig. 1E). Parental cells cultured under primed conditions remained negative throughout, whereas a discrete EGFP-positive population appeared in the reporter line by passage 1 and expanded with continued culture, comprising 15.4%, 20.9% and 28.0% of cells at passages 1, 3 and 5 respectively (Fig. 1F). The same progressive increase was observed when conversion was carried out in PXGL^18^ (Fig. S1F, G). To verify that reporter activity reflects acquisition of the naïve programme, we measured a panel of state-specific genes by RT-qPCR across the same time course. The primed marker *DUSP6* declined, while naïve genes including *NANOG*, *DPPA3*, *DNMT3L*, *KHDC1L* and *KLF17* itself increased progressively with passage (Fig. 1G). In addition, immunoblotting across primed cells and passages 1 to 7 of conversion showed that EGFP protein was undetectable in primed cells, became apparent by passage 3 and accumulated with continued culture (Fig. 1H). The reporter therefore provides a direct, antibody-free measure of naïve conversion efficiency over time.

We next compared KLF17-EGFP with established markers of the naïve state. At passage 2 of their respective primed-to-naïve conversion cultures, we sorted CD75-positive^17^ or SUSD2-positive^18^ fractions from unmodified iPSCs, EGFP-positive cells from WIBR3 ΔPE-OCT4-EGFP hESCs^7^ and KLF17-EGFP-positive cells from the reporter line, and, then profiled them by RT– qPCR (Fig. 1I). All four fractions were depleted of the primed markers *DUSP6* and *CD24*, but they differed in the expression of naïve markers: KLF17-positive and SUSD2-positive cells showed the strongest induction of *NANOG, NLRP2, DPPA5, KHDC1L, DNMT3L, DPPA3 and TFCP2L1, CD75*-positive cells were intermediate, and ΔPE-OCT4-EGFP-positive cells showed the weakest. Gating on KLF17 therefore enriches for cells with an established naïve transcriptional identity at least as effectively as the surface markers in current use.

### The reporter isolates a distinct naïve transcriptional state across culture conditions

To characterize KLF17-positive and negative populations transcriptome-wide, we converted reporter cells to naïve conditions in either HENSM or PXGL, sorted EGFP-positive and EGFP-negative fractions at passage 2, and performed RNA sequencing alongside unsorted primed cells (Fig. 2A). Principal component analysis resolved the populations along two axes (Fig. 2B). PC1, accounting for 57.4% of the variance, separated primed cells from all naïve -derived samples. PC2 (23.1%) separated the sorted fractions by reporter status rather than by medium. We next examined curated gene sets across the same samples (Fig. 2C). The primed pluripotency geneset, including *CD24, PODXL, SALL2, ZIC2, DUSP6, SFRP2, THY1* and *PSIP1*, was expressed in primed cells and silenced in naïve -derived samples, most completely in the EGFP-positive fractions. The naïve state marker genes, including *ALPG, DNMT3L, SUSD2, PBX4, KHDC3L, IL6R, CD75, GATA2*, *DPPA3* and *DPPA5*, was induced upon naïve conversion and was consistently higher in EGFP-positive than EGFP-negative cells under both culture conditions. Comparison of EGFP-positive naïve cells in HENSM with primed cells identified 2,841 upregulated and 2,175 downregulated genes, recovering the expected signature of resetting with induction of *KLF4, SUSD2, DNMT3L*, DPPA3, *DPPA5, ALPG, KHDC1L* and *KLF17* itself, and loss of *THY1, CD24, SFRP2, PODXL* and *DUSP6* (Fig. 2D). Comparison of EGFP-positive with EGFP-negative cells from the same naïve cultures identified 621 upregulated and 1,104 downregulated genes, with naïve transcripts including *ALPG, KHDC1L, DPPA3, DPPA5, DNMT3L* and *SUSD2* among the most significantly enriched in the positive fraction and primed transcripts including *SFRP2, THY1* and *CD24* retained in the negative fraction.

**Figure 2.**
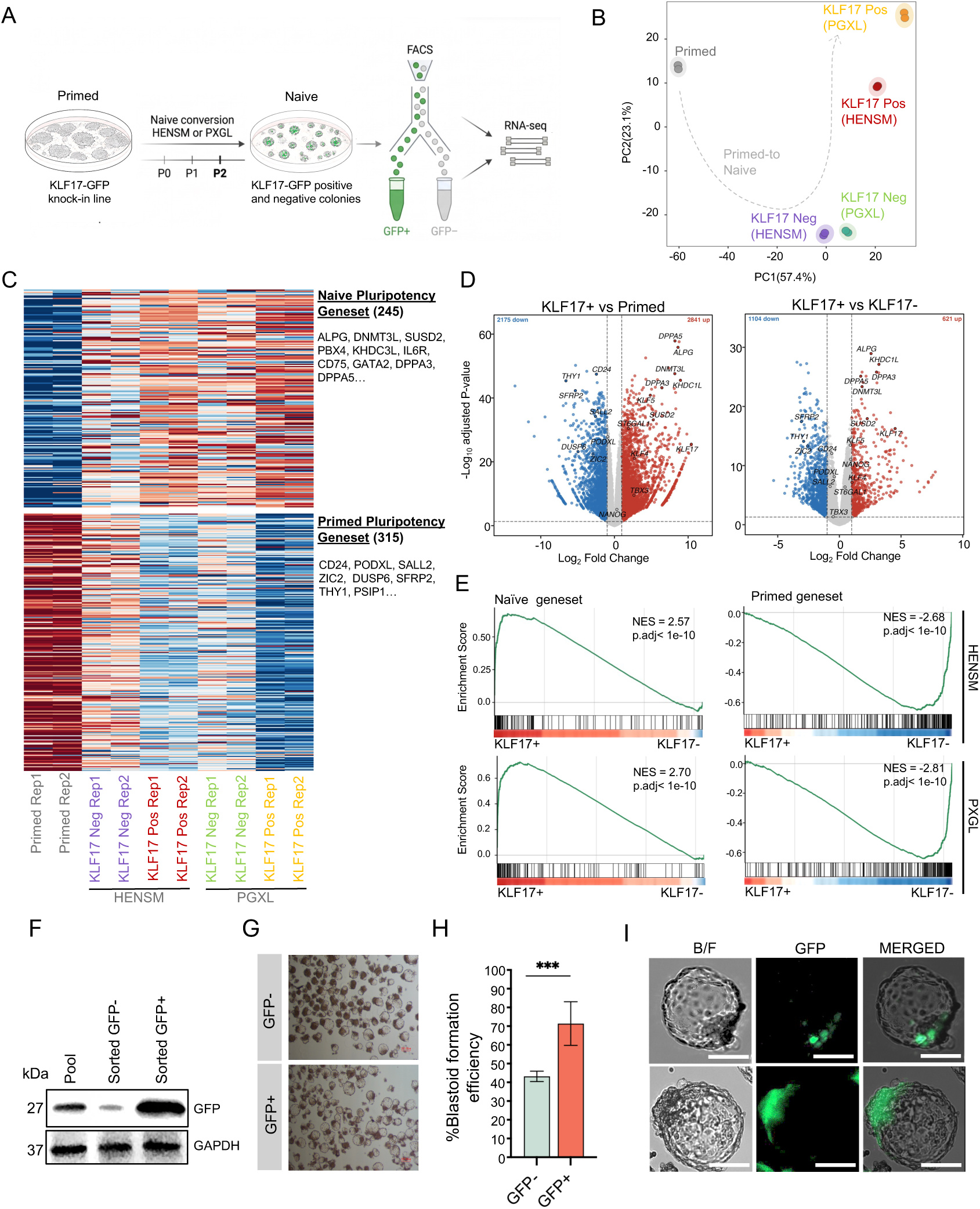
The *KLF17* reporter isolates a distinct naïve transcriptional state in two independent naïve culture conditions and marks cells competent for blastoid formation. A) Experimental scheme. KLF17-EGFP reporter cells were converted to naïve conditions in HENSM or PXGL, and EGFP-positive and EGFP-negative fractions were sorted at passage 2 for RNA sequencing. B) Principal component analysis of RNA-seq from primed cells and EGFP-positive and EGFP-negative fractions sorted from HENSM and PXGL cultures. Percentage of variance explained is indicated on each axis. Dashed line indicates the inferred primed-to-naïve trajectory. n = 2 biological replicates per group. C) Heatmap of z-scored expression across all samples for two curated gene sets: naïve pluripotency (245 genes) and primed pluripotency (315 genes). Representative genes are listed. D) Volcano plots comparing EGFP-positive naïve cells with primed cells (left) and EGFP-positive with EGFP-negative naïve cells (right), both in HENSM. Numbers of significantly up- and downregulated genes are indicated; selected genes are labelled. Thresholds: |log_2_ fold change| > 1, adjusted p < 0.05. E) Gene set enrichment analysis of the naïve (left) and primed (right) genesets in EGFP-positive versus EGFP-negative cells, in HENSM (top row) and PXGL (bottom row). NES, normalized enrichment score. F) Immunoblot for EGFP and GAPDH in the unsorted pool and in sorted EGFP-negative and EGFP-positive fractions. G) Brightfield images of the blastoids formed from EGFP-negative and EGFP-positive cells at day 5. Scale bar, 100 µm. H) Blastoid formation efficiency, scored at day 5 as the proportion of microwells containing cavitated blastocyst-like structures. Bars show mean ± SD of 2 independent experiments. ***p < 0.001. I) Brightfield, EGFP and merged images of individual blastoids. Scale bar, 100 µm.

Gene set enrichment analysis revealed that the naïve geneset was significantly enriched in EGFP-positive cells (NES = 2.57 for HENSM and 2.70 in PXGL) and the primed geneset in EGFP-negative cells (NES = −2.68 for HENSM and -2.81 in PXGL) (Fig. 2E). The same relationships were recovered when EGFP-positive PXGL cells were compared with primed cells (Fig. S2A). Sorting on the reporter alone is therefore sufficient to isolate cells that have acquired a naïve transcriptional identity from a heterogeneous resetting culture, independently of the medium used. Fold changes between the EGFP-positive and EGFP-negative fractions were highly concordant between the two media (r = 0.82 across responsive genes; Fig. S2B), and sample-to-sample correlations grouped the fractions by reporter status rather than by medium (Fig. S2C).

Because we compared two naïve media with the two reporter fractions, we could ask how much of the conversion the reporter actually marks. For each of the 15,989 expressed genes we computed the difference between EGFP-positive and EGFP-negative cells averaged over both media and the difference between the two media averaged over both fractions, and classified genes by which of the two they responded to (Log_2_FC > 1) (Fig. S2D). The largest class, 2,518 genes, had already changed in the EGFP-negative fraction relative to primed cells and changed no further with either reporter status or medium; these genes mark the response to naïve culture that both media share rather than anything the reporter distinguishes. The FGF–ERK targets *SPRY4* and *ETV1* and the primed-associated factor FOXD3 were among the most strongly downregulated genes, consistent with the MEK inhibitor PD0325901 present in both formulations, while the WNT target *SP5* together with *HES4, HOXB2, KLF5, NEAT1* and *MSX2* were among the most strongly induced. A second class of 1,097 genes differed between EGFP-positive and EGFP-negative cells, irrespective of medium used and contained well-established naïve markers, including *KLF17, KHDC1L, DPPA3, ALPG, SUSD2, CD75, DNMT3L* and *TFCP2L1*, together with the ZGA genes *TPRX1, MBD3L2B, DUXA* and *RFPL2*; on the downregulated side it contained 18 of the 33 expressed collagen genes along with *VIM, CDH2* and *ZEB2* and the primed markers *ZIC2*, *THY1* and *CRABP1*, indicating that reporter-negative cells retain a mesenchymal and extracellular-matrix programme that reporter-positive cells have extinguished (Fig. S2D). A smaller class of 177 genes responded to both variables and consisted almost entirely of the canonical ZGA repertoire, the *TRIM43, TRIM49, MBD3L* and *RFPL* families together with *ZSCAN4* and *LEUTX*, with a positive medium effect in every case, identifying this module as the source of the higher 8C-like content of naïve cultures.

We next scored each of these gene sets across all samples which reproduced these relationships (Fig. S2E). The primed set declined and the naïve set increased through the EGFP-negative to the EGFP-positive fractions in both media. The ZGA set behaved differently: it was indistinguishable between the two EGFP-negative fractions but rose sharply in EGFP-positive cells, and considerably more in HENSM than in PXGL. Gene Ontology analysis of the same comparisons placed cell adhesion, cell junction assembly and extracellular structure organization among the terms downregulated during conversion in both media, while piRNA processing and meiotic nuclear division were among those upregulated in both. Terms distinguishing HENSM from PXGL were restricted to morphogenesis and BMP-response categories, reinforcing that the two media differ far less than reporter status does (Fig. S2F). Taken together, these analyses show that both culture conditions induce a shared naïve shift, while EGFP positivity marks further acquisition of naïve and ZGA-associated programs.

Naïve hPSCs, unlike primed cells, can self-organize into blastocyst-like structures called blastoids^23–25^. These structures contain epiblast, hypoblast and trophectoderm compartments, and blastoid formation therefore provides a stringent functional assay for the naïve state. To assess whether activation of naïve transcriptional state in KLF17-positive cells correlate with higher functional potency we carried out blastoid assays using early passage sorted EGFP-positive and EGFP-negative cells which were seeded in parallel into AggreWell plates under identical blastoid formation conditions. As expected, immunoblotting for EGFP showed, enrichment of EGFP in the sorted population compared to unsorted pool (Fig. 2F). Both fractions produced cavitated, blastocyst-like structures (Fig. 2F-G), but EGFP-positive cells formed blastoids in approximately 70% of microwells compared with 40% for EGFP-negative cells (Fig. 2G, H). A single sorting step on reporter fluorescence therefore enriched for blastoid-forming competence within the same starting culture. The residual capacity of the EGFP-negative fraction is consistent with the presence of cells that had begun to reset but had not yet crossed the reporter threshold at the time of sorting. Importantly, EGFP fluorescence persisted in the resulting blastoid structures and was confined to a localized, ICM-like region of the blastoids (Fig. 2I), consistent with KLF17 expression pattern in blastocyst^26^.

### Loss of *KLF17* impairs naïve but not primed reprogramming

Although *KLF17* appears to contribute functionally to the conversion of primed PSCs to a naïve state^12,26,27^ its role in reprogramming somatic cells remains incompletely defined. Therefore, we asked whether KLF17 is required for reprogramming either to the primed or the naïve state. To test this, we targeted *KLF17* with three independent guide RNAs in human fibroblasts. Editing at each target site was confirmed by T7 endonuclease assay (Fig. 3A), and we also confirmed on target activity of the gRNAs by infecting of K2G reporter cells and observing loss of EGFP signal (Fig. 3B). To initiate reprogramming, control and knock out fibroblast were transduced with OSKM and passaged onto MEFs either under primed or naïve conditions (HENSM). Under primed conditions colony numbers were unchanged between non-targeting controls and KLF17 targeting cells (Fig. 3C). In naïve reprogramming, by contrast, all three guides significantly reduced colony formation (Fig. 3D). Naïve colonies still formed in the knockout, suggesting that KLF17 is not strictly required. These reprogramming results indicate that KLF17 is dispensable for the acquisition of primed pluripotency from somatic cells but supports efficient establishment of the naïve state.

**Figure 3.**
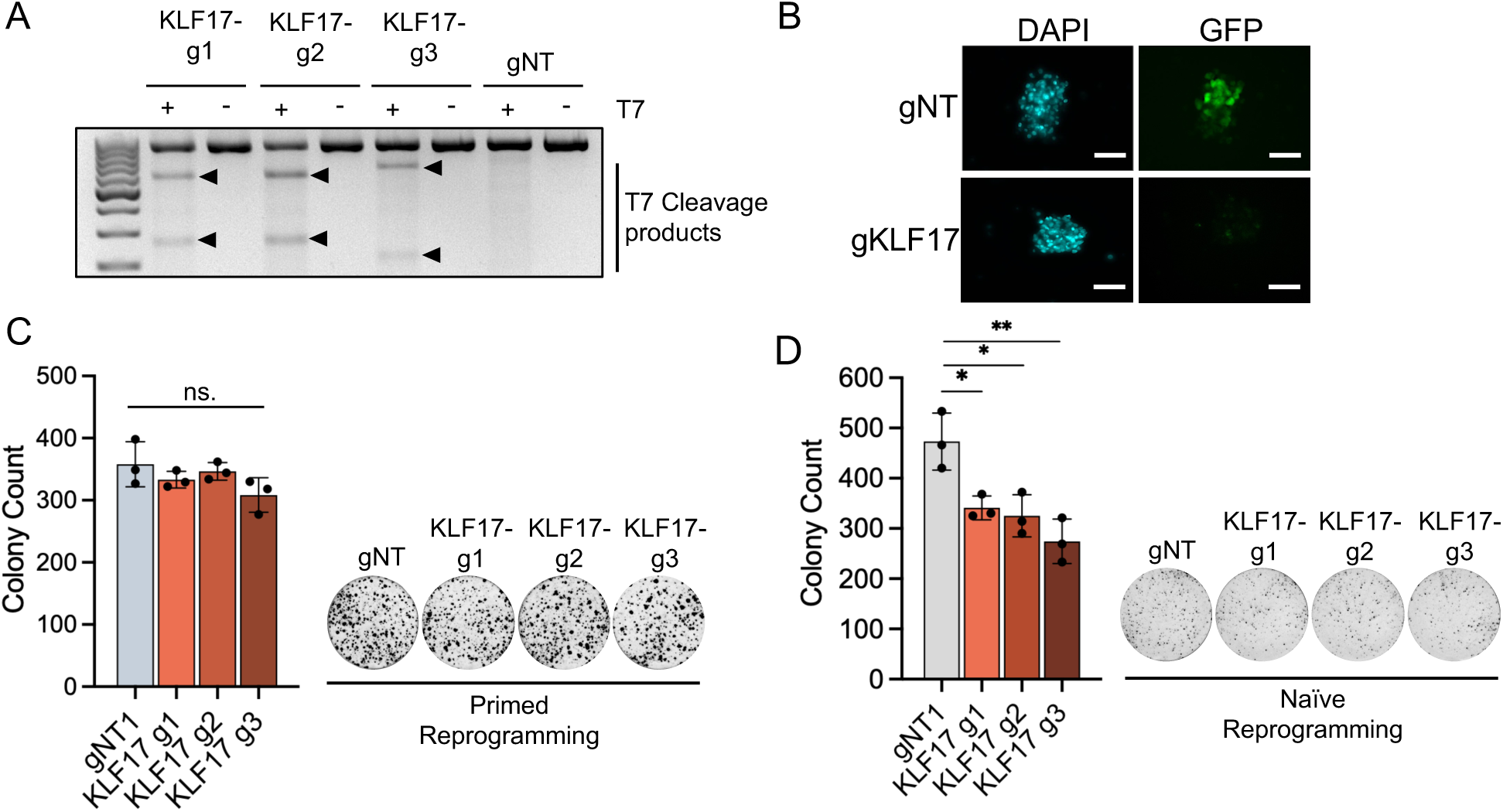
KLF17 is required for efficient naïve but not primed reprogramming. A) T7 endonuclease I assay confirming editing at each of three KLF17 target sites. gNT, non-targeting guide. Arrowheads indicate cleavage products. B) Immunofluorescence for EGFP in K2G reporter cells transduced with non-targeting or a pool of KLF17-targeting sgRNAs. Nuclei counterstained with DAPI. Scale bar, 100 µm. C) Colony counts following primed reprogramming in the presence of non-targeting or KLF17-targeting guides, with representative TRA-1-60-stained wells. Bars show mean ± SD of 3 independent experiments; individual points are shown. ns, not significant. D) As in C, for naïve reprogramming. *p < 0.05, **p < 0.01.

### *KLF17* reporter intensity resolves a graded transition toward the 8C-like state

Because *KLF17* is expressed more highly in the 8-cell embryo than in the epiblast, we asked whether higher reporter intensities would mark the naïve-to-8C-like transition. To test this directly, we treated naïve cells with TSA and DZNep to induce 8CLCs^14^ and separated the reporter positive cells into four gates; negative, low, mid and high and assessed how the transcriptome changed across them. Partitioning into four gates, untreated reporter cells distributed between the negative (52.1%) and low (40.2%) gates, with few cells reaching mid-level (3.1%) and essentially none reaching high (0.1%) (Fig. 4A). TSA/DZNep treatment redistributed the population toward higher intensities: the negative fraction fell to 29%, while mid and high fractions rose to 19.1% and 4.1% respectively (Fig. 4A, B). The scarcity of high-expressing cells in untreated naïve cultures, and their appearance only after epigenetic perturbation, is consistent with the reported frequency of spontaneous 8CLCs^13,14^ and suggested that the top gate captured a rare state. RT-qPCR-based gene expression analysis of control and TSA/DZNep-treated bulk cultures confirmed induction of naïve transcripts after treatment (Fig. S3A).

**Figure 4.**
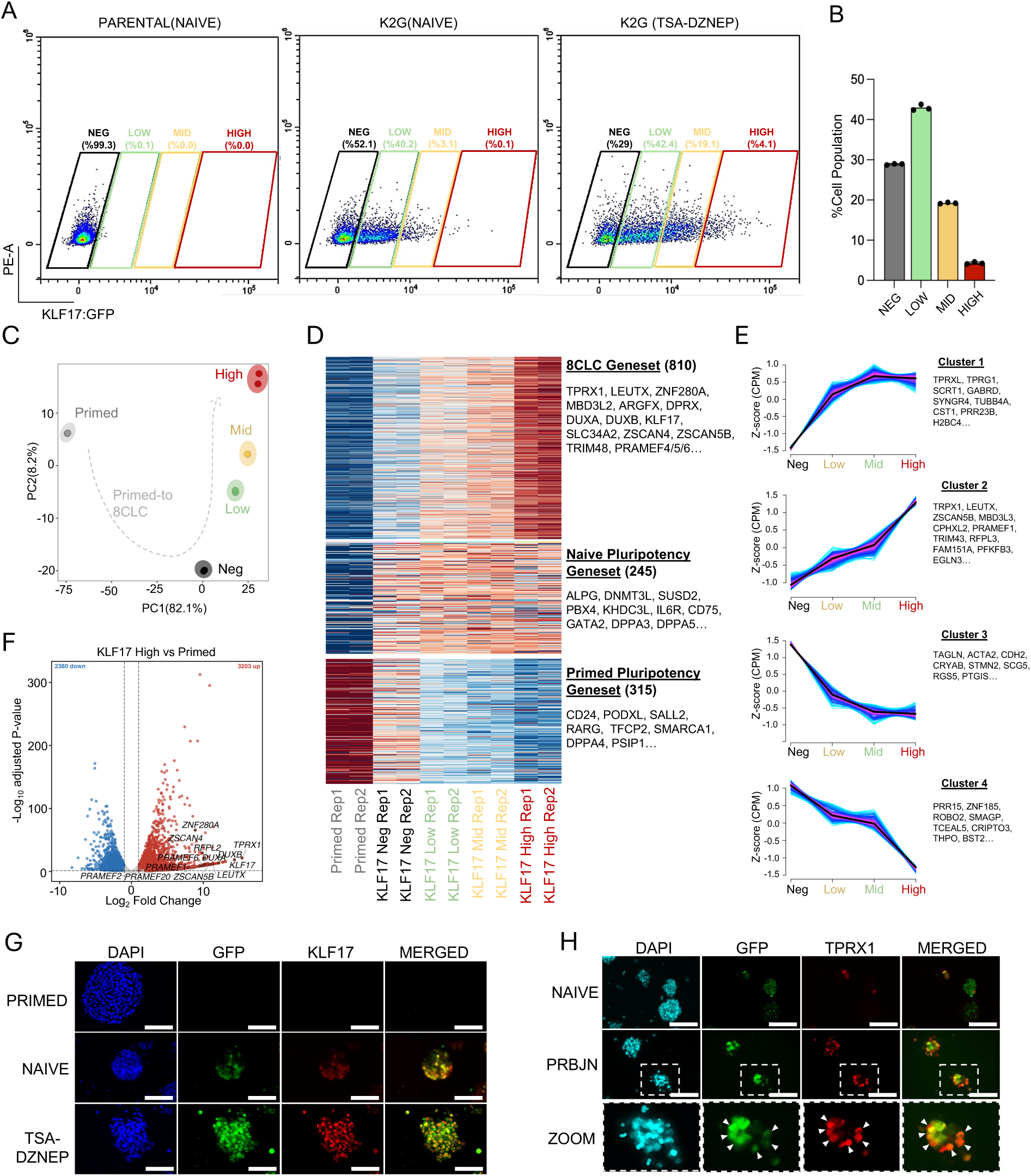
KLF17 reporter intensity resolves a graded transition toward the 8C-like state. A) Flow cytometry of the untargeted parental line under naïve conditions and KLF17:GFP reporter line under naïve and TSA/DZNep-treated conditions, resolved into negative, low, mid and high gates. Percentages indicate cells within each gate. B) Quantification of the four gates after TSA/DZNep treatment. Bars show mean ± SD. C) Principal component analysis of RNA-seq from primed cells and the four sorted gates. Percentage of variance explained is indicated on each axis. Dashed line indicates the inferred primed-to-8CLC trajectory. n = 2 biological replicates per group. D) Heatmap of z-scored expression across all samples for three curated gene sets: 8CLC (810 genes), naïve pluripotency (245 genes) and primed pluripotency (315 genes). Representative genes are listed. E) Mean z-scored expression across the four gates for the four gene clusters identified by unsupervised clustering; shading indicates the spread across genes and representative genes are listed. F) Volcano plot comparing KLF17-high cells with primed cells. Numbers of significantly up- and downregulated genes are indicated; selected genes are labelled. Thresholds: |log2 fold change| > 1, adjusted p < 0.05. G) Immunofluorescence for EGFP and KLF17 in reporter cells under primed, naïve and TSA/DZNep-treated conditions. Nuclei counterstained with DAPI. Scale bar, 100 µm. H) Immunofluorescence for EGFP and TPRX1 in naïve reporter cells and after 48 h of PRBJN treatment; the boxed region is enlarged in the lower row. Arrowheads indicate TPRX1-positive nuclei within EGFP-high cells. Nuclei counterstained with DAPI. Scale bar, 100 µm.

We sorted all four fractions from TSA/DZNep-treated cultures and performed RNA sequencing. Principal component analysis resolved every population (Fig. 4C). PC1, accounting for 82.1% of the variance, separated primed cells from all four sorted fractions. PC2 (8.2%) ordered the sorted fractions themselves, placing negative, low, mid and high samples in sequence. We next examined the expression of primed, naïve and 8CLC genes across the sorted populations (Fig. 4D). The primed pluripotency geneset was expressed in primed cells and was silenced in every naïve-derived sample, as expected. Naïve marker genes including *ALPG, DNMT3L*, *SUSD2*, *PBX4, KHDC3L, IL6R, CD75, GATA2, DPPA3* and *DPPA5* were induced upon naïve culture and retained across the sorted fractions. 8CLC genes including *TPRX1, LEUTX, ZNF280A, MBD3L2, ARGFX, DPRX, DUXA, DUXB, KLF17, SLC34A2, ZSCAN4, ZSCAN5B, TRIM48* and *PRAMEF4/5/6* increased progressively from the negative through low and mid to the high fraction, with the strongest signal confined to KLF17-high cells (Fig. 4D). This observation suggests that the reporter level enriches for the ZGA program in proportion to its intensity.

Unsupervised clustering of genes by their behavior across the four gates identified four dominant patterns (Fig. 4E). Clusters 1 and 2 comprised genes that increased with reporter intensity, differing in their kinetics: cluster 1 genes including *TPRXL*, *TPRG1*, *SCRT1*, *GABRD*, *SYNGR4*, *TUBB4A*, *CST1*, *PRR23B*, *H2BC4* rose steeply between the negative and low gates and then plateaued, whereas cluster 2 genes including *TPRX1*, *LEUTX*, *ZSCAN5B*, *MBD3L3*, *CPHXL2*, *PRAMEF1*, *TRIM43*, *RFPL3*, continued to rise across all four gates and peaked in the high fraction. The canonical ZGA transcription factors that define the 8C-like state fall in cluster 2, indicating that they are induced specifically in the KLF17 high cells rather than in any KLF17-positive cell. Clusters 3 and 4 declined with increasing intensity, the former dominated by cytoskeletal and adhesion genes (*TAGLN*, *ACTA2*, *CDH2*, *STMN2*) and the NODAL co-receptor *CRIPTO*-primed associated gene, indicating progressive suppression of epithelial-mesenchymal transition and TGF-β signaling as cells advance toward the 8C-like state.

Gene set enrichment analysis of negative vs KLF17-high sorted populations showed suppression of oxidative phosphorylation, glycolysis, PI3K-AKT-mTOR, mTORC1 and TGF-β signaling, together with epithelial-mesenchymal transition genes, and activation of the p53 pathway, TNFα/NF-κB signaling, hypoxia, the unfolded protein response, MYC targets, spermatogenesis and G2M checkpoint genes (Fig. S2G). The combination of broad metabolic suppression with induction of stress-response and cell-cycle programs is consistent with published descriptions of 8CLCs, and the enrichment of spermatogenesis-annotated genes reflects the germline-restricted expression of many ZGA factors in adult tissues rather than a germ cell identity^28^. Direct comparison of KLF17-high cells with primed cells confirmed induction of the ZGA programme, identifying 3,203 upregulated and 2,380 downregulated genes with *TPRX1*, *ZNF280A, ZSCAN4, ZSCAN5B, LEUTX, DUXA, KLF17, RFPL2* and multiple PRAMEF family members among the most significantly enriched transcripts (Fig. 4F).

We then confirmed that the reporter mirrors KLF17 expression at the protein level across both transitions. Reporter cells were maintained under primed, naïve and 8CLCs induction conditions and assessed by immunofluorescence for EGFP and endogenous KLF17 (Fig. 4G). Primed cultures showed no detectable signal for either EGFP or KLF17, confirming both that the locus is silent in this state and that the reporter contributes no background fluorescence. In naïve cultures, a subset of cells within each colony was positive for both EGFP and KLF17, with the two signals overlapping. Treatment with TSA and DZNep, two major components used to establish 8CLC induction, produced a marked increase in the intensity of both EGFP and KLF17 staining relative to untreated naïve cells.

We next asked whether the brightest reporter cells are the ones that activate the 8C program. We recently described a chemical induction protocol that can induce up to 40% cells into the 8CLC state with 48 hours^16^ . Accordingly, reporter cells were treated with PRBJN which consists of <u>P</u>Y60, <u>R</u>etinoic acid, <u>B</u>omedemstat (histone demethylase LSD1/KDM1A inhibitor), <u>J</u>QKD82 (histone demethylase KDM5 inhibitor), and <u>N</u>utlin-3a (p53 activator) and then co-stained for EGFP and TPRX1 (Fig. 4H). In control naïve cultures EGFP was present at moderate levels and TPRX1 was largely absent, whereas after PRBJN treatment TPRX1-positive nuclei appeared and were confined to the cells carrying the highest EGFP signal with the two markers overlapping. Reporter intensity therefore identifies the 8C-like subpopulation on a per cell basis.

We utilized the reporter to provide a live-cell readout for comparing 8C-inducing conditions. Naïve reporter cells were treated for 48 h with DMSO, the spliceosome inhibitor pladienolide B^29^ , TSA/DZNep or PRBJN, and the four gates were scored by flow cytometry (Fig. S3B). Each inducer shifted cells out of the negative gate, but to different extents: the KLF17-high fraction rose from 2.3% to 5.4% with pladienolide B and 4.2% with TSA/DZNep, and to 11.3% with PRBJN (Fig. S3C). The reporter therefore could rank induction protocols by how far they drive the 8CLC transition, in a single measurement on unfixed cells and without antibody staining.

As PRBJN was the most effective of the four conditions in inducing the KLF17*-*high state, we used it to ask whether KLF17 contributes functionally to the 8C-like transition rather than only reporting it. We transduced naïve reporter cells with *KLF17* targeting gRNAs and treated them with PRBJN alongside non-targeting controls. As expected EGFP signal was absent in knockout cells, but under PRBJN treatment induction of 8CLC markers such as *TPRX1, LEUTX* and *DUX4* was unaffected when compared to control cells as assessed by immunofluorescence (Fig. S3D-F). At the RNA level similar results were obtained with PRBJN induced *TPRX1, ZSCAN4, DUXB*, *KLF17, LEUTX, DUX4, ZNF280A* and *DPPA3* being at similar levels in non-targeting and KLF17-targeted cells (Fig. S3G). High KLF17 expression therefore marks the 8C-like state without being required for its induction. Taken together, these results show that a single endogenous KLF17-EGFP allele reports on two consecutive transitions in early human pluripotency.

## DISCUSSION

KLF17 has generally been studied as a marker of human naïve pluripotency, where its expression distinguishes the preimplantation epiblast and naïve hPSCs from primed cells. It was originally identified among the transcription factors enriched in the human epiblast but absent from the corresponding mouse lineage, highlighting divergence in the networks supporting pluripotency between the two species^20^. KLF17 protein becomes progressively restricted to NANOG- and SOX2-positive epiblast cells during blastocyst development^26^. Its expression during early development is nonetheless broader than this classification implies, with embryo transcriptomes placing the first major increase around the 8-cell stage, well before the epiblast. We therefore asked whether the level of KLF17 expression, rather than its presence alone, could distinguish successive developmental states in vitro. Our results show that endogenous *KLF17* spans a range in which intermediate reporter activity marks acquisition of a naïve transcriptional state, whereas substantially higher expression accompanies activation of an 8C/ZGA-associated programme.

This property distinguishes KLF17 from existing reporters of either state. ΔPE-OCT4-EGFP reports the regulatory configuration of the OCT4 locus and has been widely used to separate naïve from primed cells^7^, while surface markers such as *SUSD2* and *CD75* permit isolation without genetic engineering^17,18^. In our resetting cultures, KLF17-positive cells were enriched for naïve transcripts, but *KLF17* expression level remained informative after naïve identity had been acquired. Conversely, reporters built around TPRX1 or other components of the 8C-like programme become informative at the later transition but do not report the preceding acquisition of naïve pluripotency. The advantage of KLF17 is therefore not greater specificity for either state individually, but the ability to resolve both transitions at a single endogenous locus.

The transcriptional differences between reporter-positive and reporter-negative cells further indicate that exposure to naïve culture conditions does not produce a uniform state. Although both fractions were exposed to the same conditions, their transcriptomes were separated more strongly by KLF17 status than by whether cells were maintained in HENSM or PXGL. Reporter-negative cells had already acquired some responses to naïve culture, including repression of FGF–ERK-responsive genes, but retained higher expression of primed and extracellular-matrix-associated genes. KLF17 activation therefore marks a more advanced stage of resetting rather than the earliest response to naïve culture, consistent with previous studies identifying intermediate states during human naïve resetting^17^ .

The functional experiments suggest that KLF17 is not simply a passive marker of this transition and may help reconcile apparently different conclusions regarding its role in naïve pluripotency. Lea et al. showed that ectopic KLF17 promotes a naïve-like programme and facilitates resetting, yet KLF17-null hESCs could still acquire and maintain naïve pluripotency, with relatively modest transcriptional effects and compensatory induction of *KLF5* and *KLF4*^26^ Wang et al. reported a stronger requirement for KLF17 in the maintenance of the naïve state and identified *MAPK3* and *ZIC2* as direct targets^27^. Our results suggest that the requirement for KLF17 may depend on the route by which the naïve state is established. Targeting KLF17 did not prevent pluripotency acquisition, but colony formation was selectively reduced when fibroblasts were reprogrammed directly into naïve conditions, whereas primed reprogramming was unaffected. KLF17 therefore appears to facilitate efficient establishment of the naïve state without being absolutely required for it. This distinction may be relevant when comparing resetting of established hPSCs, maintenance of already naïve cells, and direct reprogramming from somatic cells, which begin from different regulatory landscapes. Functional redundancy among KLF factors could account for the incomplete phenotype, since *KLF4* and *KLF5* increase alongside KLF17 in our data. Separately, the proposed repression of *ZIC2* is consistent with our observations, as *ZIC2* is strongly reduced as cells enter the KLF17-positive fraction. Combinatorial perturbation of *KLF17* with related KLF factors will be required to determine whether compensation explains the residual capacity to establish naïve pluripotency.

The blastoid experiments provide an independent functional correlate of the reporter-defined naïve state. Appropriately conditioned naïve hPSCs generate blastoids at efficiencies exceeding 70%, but naïve cultures are themselves heterogeneous and the contribution of this heterogeneity to blastoid-forming competence has not been directly tested^23^. KLF17-positive cells generated blastocyst-like structures more efficiently than KLF17-negative cells isolated from the same resetting culture, linking heterogeneity in the starting population to subsequent differences in competence. However, the reporter-negative fraction retained substantial blastoid-forming capacity, and the transcriptional data instead suggest that KLF17-positive cells represent a more completely reset population at the time of sorting. Interestingly, our reporter marked ICM-like populations in blastoids, establishing it as a useful tool to monitor dynamics of epiblast identity in blastoid models.

We observed high KLF17 expression captures several central features of an 8C-like state. Canonical ZGA-associated genes, including *TPRX1, LEUTX, ZSCAN4, DUXA*, and members of the *MBD3L* and RFPL families, increase toward the KLF17-high fraction rather than uniformly across all KLF17-positive cells, as do the H3.Y-encoding genes *H3Y1* and *H3Y2*. The KLF17-high population is rare in untreated naïve cultures, at a frequency comparable to the 1.5–1.6% reported for 8CLCs by single-cell analysis and expands following treatments that promote ZGA-like transcription, while TPRX1-positive cells generated by PRBJN, a chemical cocktail we recently described for rapid 8CLC induction^16^, are concentrated among cells with the strongest reporter signal. Thus, the upper range of *KLF17* expression is associated with activation of a broader ZGA programme rather than simply stronger expression of a naïve marker.

KLF17 intensity should nevertheless not be treated as a categorical definition of an 8CLC. Previously described 8CLCs are defined by combinations of transcriptomic, transposable-element and protein-level features rather than by a single marker, and one notable difference in our data is the behaviour of *SOX2*. Taubenschmid-Stowers et al. reported reduced *SOX2* at both RNA and protein levels, whereas *SOX2* transcripts remain comparatively stable across our KLF17 intensity fractions despite strong induction of the ZGA module. KLF17-high cells may therefore encompass a broader range of ZGA-active states, including cells that have activated part of the programme while retaining features of naïve pluripotency. Differences between spontaneous and chemically induced 8CLCs may also contribute, as 8C-like populations generated by different approaches show appreciable transcriptional variation when compared with embryo references^30^.. Single-cell profiling across the KLF17 intensity ranges, would help resolve how the reporter-defined populations relate to existing 8CLC definitions.

Importantly, the strong association between high *KLF17* expression and the 8C/ZGA programme does not imply that KLF17 drives acquisition of this state. Our perturbation experiments support this distinction. Whereas KLF17 loss reduced the efficiency of naïve reprogramming, KLF17 targeting did not substantially alter the induction of *TPRX1, LEUTX*, *DUX4* and other 8C-associated markers following PRBJN treatment. This is relevant because *KLF17*-binding motifs are enriched around genes in the published 8CLC signature^13^. Our results instead fit an emerging model in which the 8C-like program can be accessed through several regulatory routes, including *DUX4*-dependent transcription, *TPRX1/DPPA3*-associated reprogramming, the *NELFA*–*TP53* axis^31^, and relief of *OTX2*-mediated repression^32^. *KLF17* may therefore lie downstream of, or respond in parallel to, these regulatory inputs rather than constituting the switch itself. Nevertheless, our results show that endogenous *KLF17* expression can be used to monitor and prospectively isolate cellular states associated with distinct stages of human preimplantation development.

## METHODS

### Cell culture

Primed hPSCs were maintained in mTeSR medium and cultured on Geltrex-coated tissue culture plates in a 5% CO_2_ and 5% O_2_ incubator. For naïve conversion cells were dissociated with Accutase and seeded at 30% confluency with 10 mM Y-27632. Medium was refreshed the next day with HENSM^12^ (1μM PD0325901, 2 μM XAV939, 1 μM Sotrastaurin, 1 μM CGP77675, 0.8 μM BIRB0796 and 20 ng/mL human LIF in N2B27 basal medium). For PXGL-based naïve culture, cells were first chemically reset and then transferred to PXGL medium as previously described^33^.Naïve PSCs were maintained under hypoxic conditions and passaged every 3-4 days using Trypsin. Medium was replaced daily throughout the experiments. For media comparison experiments, cells were adapted for at least three passages in HENSM^12^, PXGL or 5i/L/A^7^ naïve culture conditions before compound treatment. All cultures were routinely tested and confirmed negative for mycoplasma contamination.

### Generation of knock-in reporter cells

Two single guide RNAs (sgRNAs) targeting the 3’ end of *KLF17* exon 3, adjacent to available NGG PAM motifs, were designed and cloned into pSpCas9n(BB)-2A-Puro (PX462) (Addgene #62987) and verified by Sanger sequencing. The KLF17 donor template was constructed from T-2A-EGFP-PGK-Puro (Addgene #83344) where the Brachyury arms were replaced with KLF17 homology arms, leaving the rest of the cassette unchanged. Homology arms of 500 bp were defined immediately upstream and downstream of the insertion site at the *KLF17* stop codon and amplified from genomic DNA using primers carrying SacI and NheI sites for the left arm and AscI and FseI sites for the right arm (Supplementary Table 2). The left arm was first inserted between the SacI and NheI sites of the backbone, and the right arm was subsequently inserted between the AscI and FseI sites of the resulting intermediate. Each cloning step was verified by restriction digestion and Sanger sequencing across both junctions, and the final construct was sequenced in full before use.

The donor template and Cas9D10A nickase–sgRNA vectors were co-delivered into wild-type iPSCs by nucleofection. Cells were harvested with EDTA for 10 min to obtain a single-cell suspension, and approximately 1 × 10⁶ cells were resuspended in 110 µL Nucleofector R buffer (Invitrogen MPK10096) together with 2.5 µg of pX462 vectors carrying the *KLF17* sgRNAs (1.25 µg per guide) and 2.5 µg of the T2A-EGFP-PGK-Puro donor plasmid. 100 µL of the cell–DNA mixture was transferred into a cuvette pre-filled with E2 buffer and nucleofected on a 4D-Nucleofector system (Lonza) at 1200 V, 20 ms pulse length, one pulse. Cells were returned to culture at 37 °C and the medium was changed 16 h later. Nucleofected cells were selected with puromycin, dissociated to single cells and seeded into 96-well plates, and expanded clones were screened by PCR. To remove the PGK-Puro cassette from the validated knock-in line, cells were nucleofected with a Cre recombinase plasmid (pCAG-Cre-iBSD, Addgene #140284), dissociated with EDTA and seeded as single cells into 96-well plates. At approximately 50% confluency each clone was split into two wells, one maintained in mTeSR and the other treated with 1 µg/mL puromycin. Clones that failed to survive selection were recovered from the corresponding maintenance well and excision was verified by PCR.

### Flow cytometry and sorting

Cells were dissociated into single cell suspensions with 5 min trypsin incubation and resuspended in %3 FBS containing 0.5 mM EDTA/PBS. Cells were analyzed in Cytoflex S (Beckman Coulter) flow cytometer. Data were analyzed using CytExpert software. For cell sorting experiments, cells were dissociated into single cell suspensions with 5 min trypsin incubation. Pellets were resuspended in 3% FBS and 10 μM Y-27632 containing 0.5 mM EDTA/PBS, filtered using 40 μm strainers and sorted on a BD FACSAria III instrument.

### T7 endonuclease I assay

LentiCRISPRv2 plasmids carrying *KLF17-*targeting guide RNAs (1 µg) were transfected into HEK293T cells. After puromycin selection, genomic DNA was extracted using the NucleoSpin Tissue kit (Macherey-Nagel, Germany) according to the manufacturer’s instructions. The genomic region surrounding the CRISPR/Cas9 target site was amplified with Phusion high-fidelity DNA polymerase and purified using the NucleoSpin Gel and PCR Clean-up kit (Macherey-Nagel, Germany). 400 ng of purified PCR product was combined with 2 µL 10× NEBuffer 2 (NEB B7002S) and nuclease-free water to a final volume of 19 µL. Heteroduplexes were formed by denaturing at 95 °C for 5 min, ramping from 95 °C to 85 °C at −2 °C/s and from 85 °C to 25 °C at −0.1 °C/s, followed by a hold at 4 °C. 1 µL of T7 endonuclease I (NEB M0302L) was added and the reaction incubated at 37 °C for 45 min. The reaction was stopped with 1.5 µL of 0.25 M EDTA, 6 µL of 30% glycerol loading dye was added, and the products were resolved on a 2% agarose gel.

### *KLF17* Knockout Experiments

sgRNAs targeting *KLF17* were cloned into the lentiCRISPRv2 vector (Addgene #52961). Lentiviral particles were produced in HEK293T cells. For each construct, the transfer plasmid was combined with the packaging plasmid psPAX2 (Addgene #12260) and the envelope plasmid pMD2.G (VSV-G; Addgene #8454), and the plasmid mixture was added to polyethyleneimine (PEI) at a 4:1 PEI-to-DNA mass ratio in serum-free medium. After incubation at room temperature, the complexes were added dropwise to HEK293T cells. Viral supernatants were collected 48 and 72 h after transfection, cleared of cellular debris by centrifugation and filtration through a 0.45 µm filter, and concentrated by addition of 5× PEG solution.

Naïve *KLF17*-EGFP reporter iPSCs were dissociated and seeded at 1 × 10⁵ cells per well, then infected with concentrated virus in the presence of protamine sulfate (8 µg/mL). Twenty-four hours after infection and puromycin (1 µg/mL) treatment was started. Selection was continued until untransduced control wells were completely cleared, which typically took 3–4 days.

### RNA isolation and quantitative RT–PCR

Total RNA was isolated using the NucleoSpin RNA kit (Macherey-Nagel, Germany) according to the manufacturer’s instructions. cDNA was synthesized using the iScript cDNA Synthesis Kit (Bio-Rad, USA), and quantitative PCR was performed using SyberGreen mix (Takara Bio, Japan) on a LightCycler 480 real-time PCR system (Roche, Switzerland). Primer sequences are provided in Supplementary Table 2.

### Bulk RNA sequencing and analysis

Cells were sorted into the indicated fractions on a BD FACSAria III instrument. Sorted cells were pelleted by centrifugation, resuspended in DNA/RNA Shield (Zymo Research, USA) and shipped to Plasmidsaurus (USA), where RNA extraction, library preparation and sequencing were performed on an Illumina NovaSeq X Plus platform.

Raw sequencing reads were trimmed using Trimmomatic and quantified against the human reference transcriptome (GRCh38, GENCODE v48 annotation) using Salmon in quasi-mapping mode. Transcript-level estimates were imported into R using the tximport package and summarized to gene level. Differential expression analysis was performed using DESeq2.

### Somatic cell reprogramming

For knockout experiments, dH1f cells were transduced with lentiCRISPRv2 carrying *KLF17*-targeting or non-targeting sgRNAs and selected with puromycin. For reprogramming, fibroblasts were seeded at 50,000 cells per well in 12-well plates. Reprogramming factors were delivered the following day using the following lentiviruses: pSIN4-EF2-O2S (Addgene #21162) and pSIN4-CMV-K2M (Addgene #21164), encoding OCT4, SOX2, KLF4 and c-MYC. After overnight transduction the medium was replaced with fresh medium which was refreshed on days 3 and 5. For primed reprogramming, one-eighth of the cells were transferred on day 6 onto vitronectin-coated plates (Gibco A31804). From the following day the medium was switched to E7, which contains all components of E8 except TGF-β1, and was changed every 2 days until day 14 and daily thereafter until day 18. Cultures were maintained at 37°C and 5% CO₂. For naïve reprogramming, one-eighth of the cells were transferred on day 6 onto irradiated mouse embryonic fibroblast feeders. From the following day the medium was switched to HENSM and refreshed every 2 days until day 14. Cultures were maintained at 37°C, 5% O₂ and 5% CO₂.

### TRA-1-60 staining and colony quantification

Primed cultures were fixed on day 18 and naïve cultures on day 14 in 4% paraformaldehyde for 15–30 min at room temperature, then washed with PBS. For colony quantification, cells were incubated overnight at 4°C with biotinylated anti-TRA-1-60 (1:250) in PBS containing 3% fetal bovine serum and 0.3% Triton X-100, followed by streptavidin-HRP (1:500) in the same buffer for 2 h at room temperature. Colonies were visualized with DAB substrate and TRA-1-60-positive colonies were counted in ImageJ.

### Western blotting

Cells were lysed in RIPA buffer supplemented with protease inhibitor cocktail. Equal amounts of protein were separated by SDS–PAGE and transferred onto PVDF membranes. Membranes were blocked with 5% non-fat milk in PBST for 1 h at room temperature and incubated overnight at 4°C with primary antibodies against KLF17 (1:1,000), EGFP (1:2,000) and GAPDH(1:2000). After washing with PBST, membranes were incubated with the appropriate HRP-conjugated secondary antibodies. Protein bands were detected using enhanced chemiluminescence (ECL) reagents according to the manufacturer’s instructions.

### Immunofluorescence staining

Cells were fixed in 4% paraformaldehyde for 15 min at room temperature (RT) and washed three times with PBS. Cells were then permeabilized with 0.5% Triton X-100 in PBS for 10 min and blocked in PBS containing 3% serum for 1 hr at RT. Primary antibodies (Anti-KLF17 (Sigma, HPA024629) 1:300, Anti-LEUTX (Novus Biologicals, NBP1-90890) 1ug/ml and Anti-DUX4 (Sigma, SAB5200019) 1:300 were incubated overnight at 4°C followed by fluorophore-conjugated secondary antibodies (Goat anti-Rabbit IgG Alexa Fluor 488 (Thermo Fisher, cat# A11034)). Nuclei were counterstained with 1 µg/ml DAPI for 5 min. Images were acquired using LEICA DMI8 confocal microscope and processed using ImageJ.

### Induction of 8C-like cells

For TSA/DZNep induction, naïve cells were treated with trichostatin A (20 nM) and DZNep (50 nM) for 48 h. For PRBJN induction, naïve cells were treated for 48 h with a five-component cocktail consisting of PY60 (7.5 µM), Retinoic acid (0.5 µM), Bomedemstat (1 µM), JQKD82 (1 µM) and Nutlin-3a (0.5 µM), as described^16^. Control cultures received 0.1% DMSO. All treatments were performed under hypoxic conditions (5% O₂, 5% CO₂).

### Induction of Blastoids

Blastoid generation experiments were approved by the Koç University Biomedical Research Ethics Committee (2025.068.IRB2.031) and followed The International Society for Stem Cell Research (ISSCR) Guidelines for Stem Cell Research and Clinical Translation^34^. KLF17-EGFP reporter cells maintained in HENSM on Geltrex-coated plates were dissociated using 1 mM EDTA in PBS for 10–15 min at room temperature, and EGFP-positive and EGFP-negative fractions were isolated by fluorescence-activated cell sorting. Sorted cells were collected by centrifugation and resuspended in aggregation medium consisting of N2B27 supplemented with 0.3% BSA and 1× CEPT cocktail. AggreWell 400 plates were prepared by treatment with Anti-Adherence Rinsing Solution, followed by centrifugation at 500 × g for 1 min, sonication for 2 min to remove trapped air bubbles, and incubation at room temperature for at least 1 h. Wells were then washed twice with N2B27 medium and seeded into AggreWell 400 plates at a density of 35 cells per microwell. Plates were incubated at room temperature for 15 min and centrifuged at 300 × g for 3 min. Aggregates were cultured in aggregation medium supplemented with 1× CEPT cocktail for the first 24 hours after which, half of the medium was replaced with 2× PALLY medium consisting of N2B27 supplemented with PD0325901 (2 μM), A83-01 (2 μM), lysophosphatidic acid (LPA; 5 μM), human LIF (80 ng/mL), and 1× CEPT cocktail. On day 2, the medium was completely replaced with 1× PALLY medium. On days 3 and 4, cultures were maintained in N2B27 supplemented with LPA (2.5 μM), and 1× CEPT cocktail. Thereafter, the medium was refreshed every 24 h with the same formulation for the remainder of the culture period. All cultures were maintained under hypoxic conditions (5% O₂ and 5% CO₂) throughout blastoid generation.

## Data availability

All sequencing data generated in this study have been deposited in the Gene Expression Omnibus (GEO) under the accession number GSE350630.

## Author contributions

A.O, S.U and T.T.O. conceived ideas; N.K and S.U generated KLF17-reporter line. S.U and A.O performed naïve conversion, blastoid generation, flow cytometry, immunofluorescence staining, RNA sequencing. A.O performed bioinformatic analyses; A.O and T.T.O. wrote the manuscript.

## Materials & Correspondence

The plasmid generated in this study is available from the corresponding author upon reasonable request.

## Acknowledgment

We thank Koç University School of Medicine Department of Medical Genetics for karyotyping. The authors gratefully acknowledge the use of the services and facilities of the Koç University Research Center for Translational Medicine (KUTTAM).

---

**Supplementary Figure 1.**
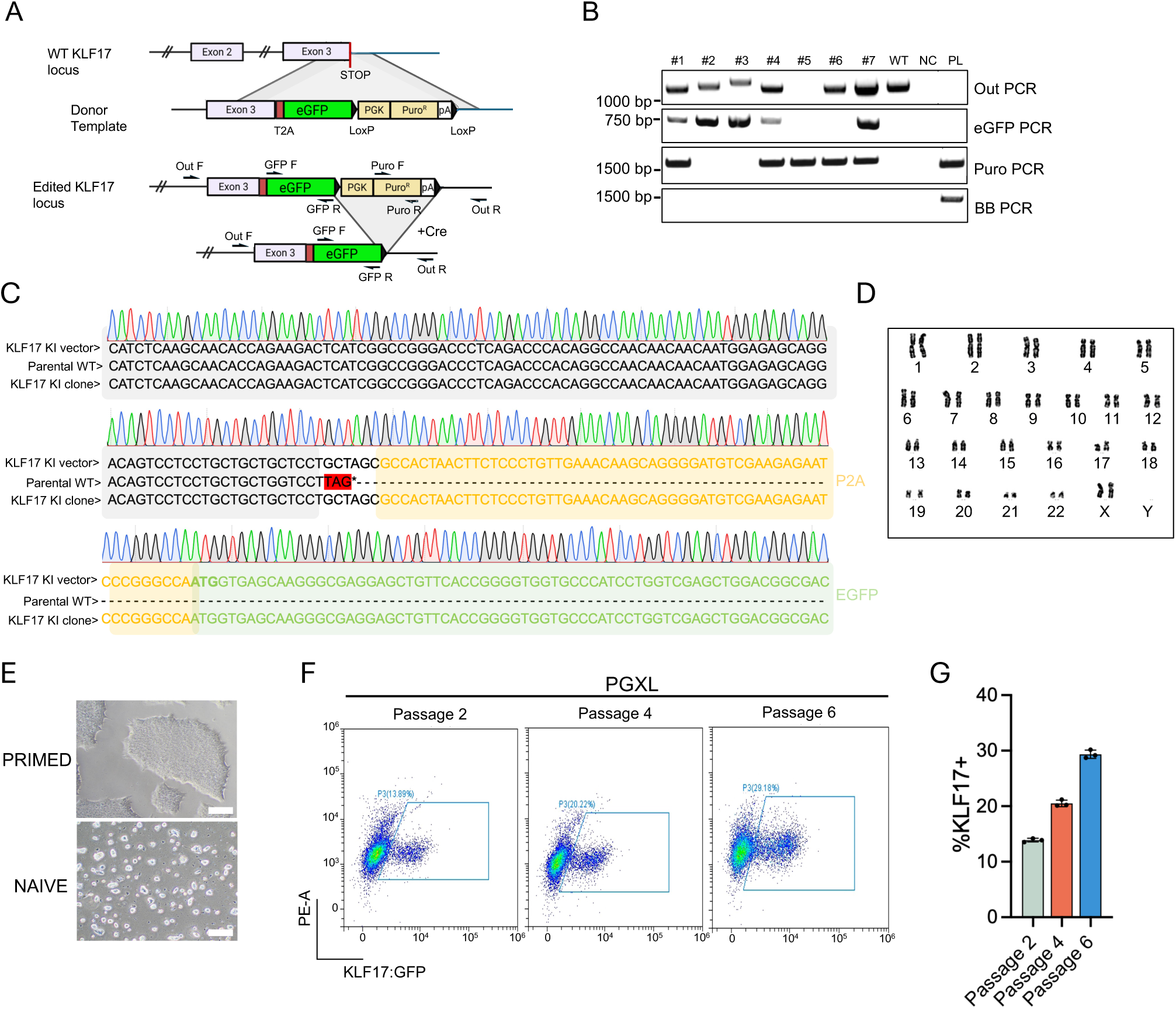
A) Targeting strategy for generating EGFP knock-in at the KLF17 locus. A T2A-EGFP cassette followed by a LoxP-flanked PGK-Puro selection cassette was inserted into the last exon of KLF17 immediately upstream of the stop codon. Arrows indicate primers used for genotyping; the bottom schematic shows the locus after Cre-mediated excision of the LoxP-flanked selection cassette. B) Genotyping PCR across candidate clones using the indicated primer sets. Out PCR spans the targeted region; EGFP and Puro PCRs detect the inserted cassettes; BB PCR detects donor plasmid backbone and controls for random integration. WT, parental line; NC, no-template control; PL, donor plasmid. C) Sanger sequencing of the edited locus in the selected clone, aligned to the donor vector and the parental wild-type sequence, showing in-frame insertion of T2A-EGFP in place of the stop codon. D) G-banded karyotype of the reporter line after targeting. E) Phase-contrast images of the reporter line in primed and naïve conditions. Scale bar, 100 µm. F) Flow cytometry of KLF17:EGFP in the reporter line at passages 2, 4 and 6 of naïve conversion in PXGL medium. G) Percentage of EGFP-positive cells in primed cells and at passage 6 of naïve conversion in PXGL. Bars show mean ± SD of 3 experiments. ****p < 0.0001.

**Supplementary Figure 2.**
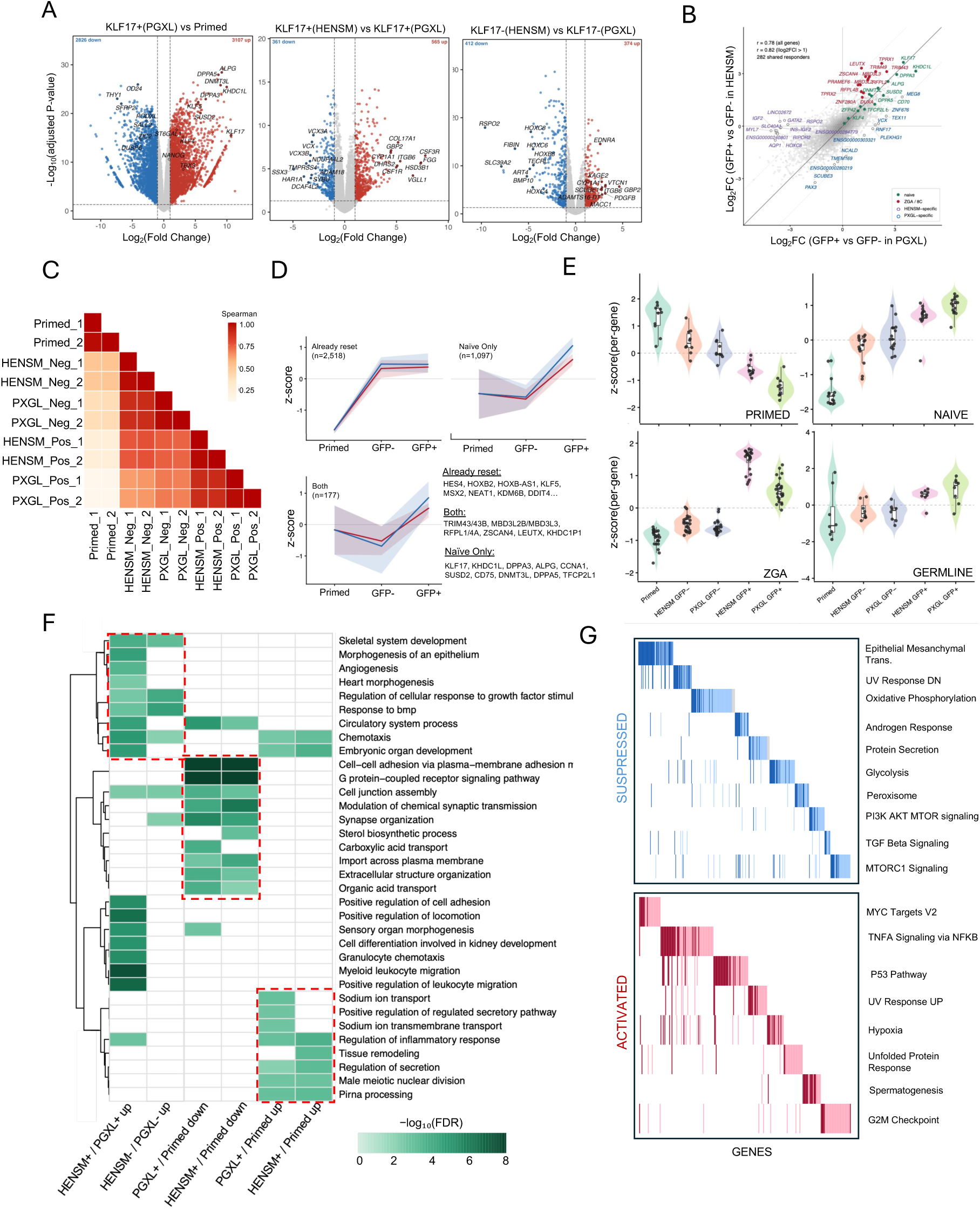
A) Volcano plots comparing EGFP-positive PXGL cells with primed cells (left), EGFP-positive HENSM with EGFP-positive PXGL cells (middle), and EGFP-negative HENSM with EGFP-negative PXGL cells (right). Numbers of significantly up- and downregulated genes are indicated; selected genes are labelled. B) Log2 fold change between EGFP-positive and EGFP-negative fractions in HENSM plotted against the same comparison in PXGL. Each point is one gene; naïve and 8C-associated genes are highlighted, as are genes responding in only one medium. The diagonal indicates an identical response in the two media; r values for all genes and for genes responding in at least one medium (|log2FC| > 1) are indicated. C) Spearman correlation between samples, calculated on the 2,000 most variable genes. D) Genes classified by whether they respond to reporter status, to the medium, or to neither, shown as the mean z-score across primed, EGFP-negative and EGFP-positive samples; the three largest classes are shown, shading indicates the interquartile range and representative genes are listed. E) Per-gene z-scores of the primed, naïve, 8C-associated and germline gene sets across all samples; each point is one gene, violins show the distribution and boxes the interquartile range. F) Gene Ontology biological process terms enriched among the genes up- or downregulated in each comparison, colored by −log10 FDR and capped at 8. G) Gene set enrichment analysis of KLF17-high versus KLF17-negative cells, showing significantly suppressed (blue) and activated (red) hallmark gene sets. Each vertical line represents one gene.

**Supplementary Figure 3.**
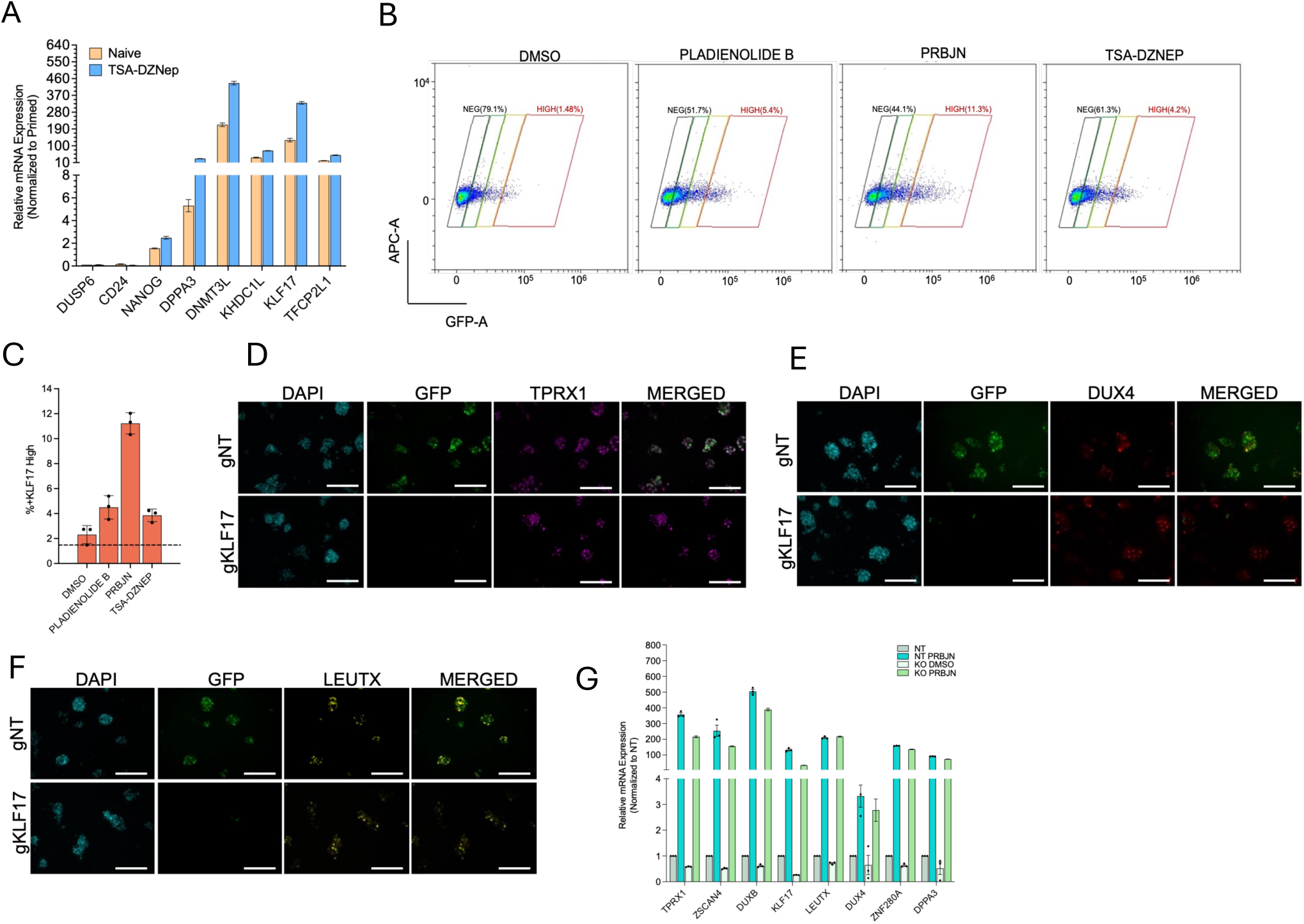
A) RT-qPCR for primed-, naïve- and 8C-associated genes in naïve and TSA/DZNep-treated cells. Expression normalized to GAPDH and shown relative to primed. Mean ± SD of 3 biological replicates. B) Flow cytometry of KLF17:EGFP in naïve reporter cells after 48 h of treatment with DMSO, pladienolide B, PRBJN or TSA/DZNep, resolved into negative, low, mid and high gates. C) Quantification of the KLF17-high fraction across the four conditions. Bars show mean ± SD of 3 independent experiments; individual points are shown. Dashed line, DMSO baseline. D) Immunofluorescence for EGFP and TPRX1 in non-targeting (gNT) and KLF17-targeted (gKLF17) naïve reporter cells after 48 h of PRBJN treatment. Nuclei counterstained with DAPI. Scale bar, 200 µm. E) As in D, for DUX4. F) As in D, for LEUTX. G) RT-qPCR for 8C-associated genes in non-targeting (NT) and KLF17-knockout (KO) cells under control (DMSO) and 8CLC-inducing (PRBJN) conditions. Expression normalized to GAPDH and shown relative to NT. Mean ± SD of 3 technical replicates.

## References

1. Rossant, J. & Tam, P. P. L. New Insights into Early Human Development: Lessons for Stem Cell Derivation and Differentiation. Cell Stem Cell 20, 18–28 (2017).

2. Thomson, J. A. et al. Embryonic Stem Cell Lines Derived from Human Blastocysts. Science 282, 1145–1147 (1998).

3. O’Leary, T. et al. Tracking the progression of the human inner cell mass during embryonic stem cell derivation. Nat. Biotechnol. 30, 278–282 (2012).

4. Takahashi, K. et al. Induction of Pluripotent Stem Cells from Adult Human Fibroblasts by Defined Factors. Cell 131, 861–872 (2007).

5. Nakamura, T. et al. A developmental coordinate of pluripotency among mice, monkeys and humans. Nature 537, 57–62 (2016).

6. Takashima, Y. et al. Resetting Transcription Factor Control Circuitry toward Ground-State Pluripotency in Human. Cell 158, 1254–1269 (2014).

7. Theunissen, T. W. et al. Systematic Identification of Culture Conditions for Induction and Maintenance of Naive Human Pluripotency. Cell Stem Cell 15, 471–487 (2014).

8. Guo, G. et al. Epigenetic resetting of human pluripotency. Development 144, 2748–2763 (2017).

9. Guo, G. et al. Naive Pluripotent Stem Cells Derived Directly from Isolated Cells of the Human Inner Cell Mass. Stem Cell Rep. 6, 437–446 (2016).

10. Liu, X. et al. Comprehensive characterization of distinct states of human naive pluripotency generated by reprogramming. Nat. Methods 14, 1055–1062 (2017).

11. Kilens, S. et al. Parallel derivation of isogenic human primed and naive induced pluripotent stem cells. Nat. Commun. 9, 360 (2018).

12. Bayerl, J. et al. Principles of signaling pathway modulation for enhancing human naive pluripotency induction. Cell Stem Cell 28, 1549–1565.e12 (2021).

13. Taubenschmid-Stowers, J. et al. 8C-like cells capture the human zygotic genome activation program *in vitro*. Cell Stem Cell 29, 449–459.e6 (2022).

14. Mazid, M. A. et al. Rolling back human pluripotent stem cells to an eight-cell embryo-like stage. Nature 605, 315–324 (2022).

15. Yu, X. et al. Recapitulating early human development with 8C-like cells. Cell Rep. 39, 110994 (2022).

16. Odabas, A. et al. Rapid and efficient generation of human 8-cell-like cells for embryo modelling. 2026.08.18.745473 Preprint at 10.64898/2026.08.18.745473 (2026).

17. Collier, A. J. et al. Comprehensive Cell Surface Protein Profiling Identifies Specific Markers of Human Naive and Primed Pluripotent States. Cell Stem Cell 20, 874–890.e7 (2017).

18. Bredenkamp, N., Stirparo, G. G., Nichols, J., Smith, A. & Guo, G. The Cell-Surface Marker Sushi Containing Domain 2 Facilitates Establishment of Human Naive Pluripotent Stem Cells. Stem Cell Rep. 12, 1212–1222 (2019).

19. Yoshihara, M. et al. Transient DUX4 expression in human embryonic stem cells induces blastomere-like expression program that is marked by SLC34A2. Stem Cell Rep. 17, 1743– 1756 (2022).

20. Blakeley, P. et al. Defining the three cell lineages of the human blastocyst by single-cell RNA-seq. Development 142, 3151–3165 (2015).

21. Petropoulos, S. et al. Single-Cell RNA-Seq Reveals Lineage and X Chromosome Dynamics in Human Preimplantation Embryos. Cell 165, 1012–1026 (2016).

22. Stirparo, G. G. et al. Integrated analysis of single-cell embryo data yields a unified transcriptome signature for the human pre-implantation epiblast. Development 145, dev158501 (2018).

23. Kagawa, H. et al. Human blastoids model blastocyst development and implantation. Nature 601, 600–605 (2022).

24. Yanagida, A. et al. Naive stem cell blastocyst model captures human embryo lineage segregation. Cell Stem Cell 28, 1016–1022.e4 (2021).

25. Wei, Y. et al. Generation of Human Blastoids from Naive Pluripotent Stem Cells. Methods Mol. Biol. 2767, 1–18 (2024).

26. Lea, R. A. et al. KLF17 promotes human naïve pluripotency but is not required for its establishment. Development 148, dev199378 (2021).

27. Wang, S.-H., et al. KLF17 promotes human naive pluripotency through repressing MAPK3 and ZIC2. Sci. China Life Sci. 65, 1985–1997 (2022).

28. Simpson, A. J. G., Caballero, O. L., Jungbluth, A., Chen, Y.-T. & Old, L. J. Cancer/testis antigens, gametogenesis and cancer. Nat. Rev. Cancer 5, 615–625 (2005).

29. Li, S. et al. Capturing totipotency in human cells through spliceosomal repression. Cell 187, 3284–3302.e23 (2024).

30. Yoshihara, M. & Kere, J. Transcriptomic differences between human 8-cell-like cells reprogrammed with different methods. Stem Cell Rep. 18, 1621–1628 (2023).

31. Su, X. et al. NELFA supports naïve pluripotency and drives 8C-like state in human embryonic stem cells. Nat. Commun. 10.1038/s41467-026-72714-z(2026) doi:10.1038/s41467-026-72714-z.

32. Kong, X. et al. OTX2 inhibits human pluripotent stem cell reprogramming toward 8-cell-like and morula-like states. Nat. Commun. 17, 1685 (2026).

33. Rugg-Gunn, P. J. Induction of Human Naïve Pluripotency Using Chemical Resetting. Methods Mol. Biol. 2416, 29–37 (2022).

34. Clark, A. T. et al. Stem cell-based embryo models: The 2021 ISSCR stem cell guidelines revisited. Stem Cell Rep. 20, 102514 (2025).

